# Targeted phage hunting to specific *Klebsiella pneumoniae* clinical isolates is an efficient antibiotic resistance and infection control strategy

**DOI:** 10.1101/2024.01.07.574526

**Authors:** Celia Ferriol-González, Robby Concha-Eloko, Mireia Bernabéu-Gimeno, Felipe Fernández-Cuenca, Javier E. Cañada-García, Silvia García-Cobos, Rafael Sanjuán, Pilar Domingo-Calap

**Affiliations:** Instituto de Biología Integrativa de Sistemas, Universitat de València-CSIC, Paterna, Spain; Unidad Clínica de Enfermedades Infecciosas y Microbiología, Hospital Universitario Virgen Macarena, Sevilla, Spain; Instituto de Biomedicina de Sevilla, Hospital Universitario Virgen Macarena-CSIC-Universidad de Sevilla, Sevilla, Spain; CIBER de Enfermedades Infecciosas (CIBERINFEC), Instituto de Salud Carlos III, Madrid, Spain; Laboratorio de Referencia e Investigación en Resistencia a Antibióticos e Infecciones Relacionadas con la Asistencia Sanitaria, Centro Nacional de Microbiología, Instituto de Salud Carlos III, Madrid, Spain

**Keywords:** *Klebsiella pneumoniae*, phage hunting, phage therapy, phage cocktail, host range, bacterial capsule, receptor binding protein, depolymerase

## Abstract

*Klebsiella pneumoniae* is one of the most threatening multi-drug resistant pathogens today, with phage therapy being a promising alternative for personalized treatments. However, the intrinsic capsule diversity in *Klebsiella* spp. poses a substantial barrier to phage host range, complicating the development of broad-spectrum phage-based treatments. Here, we have isolated and genomically characterized phages capable of infecting each of the acquired 77 reference serotypes of *Klebsiella* spp., including capsular types widespread among high-risk *K. pneumoniae* clones causing nosocomial infections. We demonstrated the possibility of isolating phages for all capsular types in the collection, revealing high capsular specificity among taxonomically related phages, in contrast to a few phages that exhibited broad-spectrum infection capabilities. To decipher the determinants of the specificity of these phages, we focused on their receptor-binding proteins, with particular attention to depolymerase domains. We also explored the possibility of designing a broad-spectrum phage cocktail based on phages isolated in reference capsular type strains, and determining the ability to lysate relevant clinical isolates. Interestingly, a combination of 12 phages capable of infecting 60% of the reference *Klebsiella* spp. serotypes was tested on a panel of carbapenem-resistant *K. pneumoniae* clinical isolates. Our results suggest that in a highly variable encapsulated bacterial host, phage hunting must be directed to the specific *Klebsiella* isolates. This work is a step forward in the understanding of the complexity of phage-host interactions, and highlights the importance of implementing precise and phage-specific strategies to treat *K. pneumoniae* infections worldwide.

## Introduction

*Klebsiella pneumoniae,* an opportunistic Gram-negative encapsulated bacterium, belongs to the ESKAPE group that comprises the most critical multi-drug resistant (MDR) bacterial pathogens (1,2). Worldwide, more than 69,000 deaths per year are associated with MDR *K. pneumoniae,* being considered a major threat (3). Indeed, *K. pneumoniae* is considered the fastest growing pathogenic bacterium in the European region currently, with high prevalence in hospital-acquired infections (4,5). Although it typically colonizes human gastrointestinal tract asymptomatically as commensals, in immunocompromised and critically ill patients, *K. pneumoniae* can lead to severe diseases such as pneumonia, urinary tract infections, bloodstream infections, and wound or surgical site infections (6). The main virulence factor of *Klebsiella* spp. is its capsule, which protects bacteria from the immune system, but also from adverse environmental conditions and possible predators (7,8). *Klebsiella* is a highly variable bacterium, with 77 capsule serotypes (9–11), identified as the reference capsular types. However, high-throughput sequencing and bioinformatics tools have led to more than 180 capsular locus types (KL-types) described (12,13). Traditionally, 7-gene multilocus sequence typing (MLST) nomenclature has been widely used to describe *K. pneumoniae* high-risk clones (12,14). Indeed, these STs can be associated to specific KL-types (12). In Spain, the most abundant circulating in 2019 were ST307, ST11, ST512, ST15, ST147 and ST392 (13). Furthermore, many of these STs are distributed worldwide producing nosocomial outbreaks (5) and have accumulated different carbapenemases, with the most prevalent being *bla*_OXA-48_, *bla*_KPC-3_, *bla*_VIM-1_, and *bla*_NDM-1_ (13,15).

The lack of effective treatments for MDR bacteria forces the implementation of alternative strategies, among which bacteriophage (phage) therapy emerges as a promising option (16). Phages are viruses that infect bacteria, and they were used for treating bacterial infections since their early discovery (17–19). However, for therapy, only strictly lytic phages are useful, avoiding phages encoding lysogenic genes and preventing gene transfer. A key feature of phages is their specificity. Most phages are capable of infecting a small subset of strains within a bacterial species (20,21), while other phages exhibit broader host ranges, being able to infect even different bacterial species (22). Phages with narrow host range may be considered advantageous for personalized phage therapy, as it allows the phage to target pathogenic bacteria without affecting the microbiota, reducing the risk of side effects (23). However, broad host range phages also present some advantages, since they can lyse a wide range of bacterial strains, making them particularly useful when multiple bacteria are responsible for an infection, or as preventive tools (24,25).

Bacteria, as evolving entities, can develop phage-resistance mechanisms. Indeed, it is known that *Klebsiella* spp. can emerge phage-resistance in a few hours (26). Interestingly, phages can overcome this resistance, since they co-evolve with bacteria as an arms race between the two components (27). In this sense, both narrow and broad host range phages are susceptible to inducing bacterial phage resistance, and phage cocktails may overcome or delay this emergence. Developing phage cocktails able to control bacteria as preventive tools is a challenge, since there are many limitations in the phage-host interactions. However, understanding phage diversity could provide new insights in the development of targeted strategies of phage hunting and the implementation of new research lines. It is known that phage diversity is enormous, but we are far away from isolating and being able to grow and culture the vast majority of phages.

Targeted phage hunting to specific hosts could be an interesting approach to create a large collection of phages with multitude of applications. In this sense, *K. pneumoniae* is a major challenge due to its capsular type diversity, which has been proposed as the main restrictive determinant of phage host range (28). Understanding the mechanisms underlying phage host range is important to develop new treatments, being host recognition the first barrier and a major host-range determinant (28). Phage receptor binding proteins (RBP) mediate the initial phage-host recognition, joining to the receptor of the host surface (29). Bacterial capsules can mask cell receptors, protecting them from phage infection (30), although they can also be exploited by phages for its attachment (31,32). To overcome the barrier that capsule poses for the infection, some phages encode specific hydrolases called depolymerases (Dpos), enzymes able to recognize, bind and digest specifically the oligosaccharide bounds. Phage Dpos are domains of the RBPs usually known as tail spikes, and some of them are produced as soluble proteins (33,34). In addition to this first-step barrier, post-adsorptive defense mechanisms have a role in the phage infectivity, such as restriction mechanisms (RM) (35), abortive infection systems (Abi) (36), regularly interspaced short palindromic repeats (CRISPRs) (37,38) and many other defenses that have being discovered recently (39,40).

Exploring deeper the host range diversity in *Klebsiella* phages is needed to implement phage-based control strategies. Interestingly, we can create cocktails *à la carte*, depending on the epidemiology of a given region, including phages with complementary host ranges, achieving a broader spectrum for the whole preparation (41,42), or presenting synergistic effects that improve their efficacy (43,44). Here, we have isolated and characterized a myriad of phages, suggesting the feasibility to develop precise and personalized treatments based on environmental phages. The evaluation of the cross-infectivity of the phages allowed us to unravel the underlying factors determining the host range of *Klebsiella* phages, showing that they can be associated in clusters. In deep analyses of the Dpos and RBPs reported unraveled diversity and provided the basis to understand phage-host interactions. With this information, we designed a broad-spectrum phage cocktail against high-risk clones of *K. pneumoniae* circulating in Spain. Our results showed that the high specificity of *Klebsiella* phages and intrinsic bacterial defense mechanisms impair phage infectivity. Thus, phage hunting directed to the clinical strain must be a promising solution to combat *K. pneumoniae* infections. Indeed, we suggest that phage cocktails should be limited to prevention strategies or ready-to-use treatments, specifically designed for requirements in each region based on the epidemiology.

## Materials and Methods

### Bacterial strains and sequencing

The 77 *Klebsiella* spp. reference strains collection, corresponding to the *Klebsiella* spp. reference serotypes, was purchased from the Statens Serum Institut (Copenhagen, Denmark). In addition, 58 carbapenem-resistant *K. pneumoniae* clinical isolates corresponding to the high-risk clones circulating in Spain between 2019 and 2021 were selected for this study (Supplementary materials 1). They were all collected from clinical samples (urine, blood, surgical devices, and diverse swabs and exudates) in Spanish hospitals and stored by the *Programa Integral de Prevención y Control de las Infecciones Relacionadas con la Asistencia Sanitaria y Uso Apropiado de los Antimicrobianos* (PIRASOA), at the Instituto de Biomedicina de Sevilla (IBIS, Sevilla, Spain), and the CARB-ES-19 project (13), and the *Programa de Vigilancia del Laboratorio de Referencia e Investigación en Resistencia a Antibióticos e Infecciones Relacionadas con la Asistencia Sanitaria* at the Centro Nacional de Microbiología (CNM, Instituto de Salud Carlos III, Spain). Genomic DNA libraries were prepared using Nextera DNA Flex Library Preparation Kit (Illumina Inc., San Diego, CA, United States) and whole-genome sequencing was performed using Illumina NextSeq 550 according to the manufacturer’s instructions, generating paired-end (2×150) reads. Quality of the reads was assessed using FASTQC (v0.11.9) (45) before their submission to the European Nucleotide Archive (PRJEB68301). After *de nov*o genome assembly using Unicycler (v0.4.8) (46), quality was assessed with QUAST (v5.2.0) (47) and ST, K-locus (KL), O-Antigen (O-Ag) and carbapenemase-encoding genes were identified using Kleborate (v2.3.2) (12) using CARD database for the detection of antimicrobial resistance genes (48).

### Phage isolation, purification and DNA extraction

A high-throughput screening was performed for targeted phage hunting. Phages were isolated from environmental samples collected from wastewater treatment plants and surrounding areas in Valencia (Spain). Plaque isolation was done as described before (26). The isolation of phages in *K. pneumoniae* reference strain with capsule serotype 20 (K20) was performed differently due to technical issues. In this case, prior enrichment of the environmental samples was needed, incubating the samples with the host overnight (37°C, 200 rpm). After 24 hours, lysates were centrifuged twice (3,900 rpm, 5 min) to remove bacteria. For phage concentration, all phages were amplified for 3h in a final volume of 5 mL in Luria-Bertani (LB) with CaCl2 (37°C, 200 rpm), and centrifuged twice after the amplification (3,900 rpm, 5 min) to remove bacteria. Lysates were centrifuged in a high-speed centrifuge (80,000 rpm, 3 hours), and pellets were resuspended in 200 µL of SM buffer (50 mM Tris-HCl [pH 7.5], 8 mM MgSO4•7H2O, 100 mM NaCl, and 0.01% gelatin [w/v]). Removal of host DNA and digestion of phage capsids were done as described before (28). Extraction and purification of DNA was performed using DNA Clean & Concentrator 5-Kit (Zymo) or Maxwell PureFood GMO and Authentication Kit (Promega) with Maxwell RSC Instrument (Promega).

### Phage sequencing and assembly

Sequencing libraries were prepared using Illumina Nextera XT DNA kit (paired-end reads 2×250 bp or 2×150 bp), and reads were generated in the Illumina MiSeq platform with iSeq Reagent Kit v2. For potential contaminant analysis, the Kraken 2 tool was used (49), while sequencing read quality was assessed using FastQC software (version 0.11.9, Babraham Bioinformatics) (45). *De novo* genome assembly was carried out using either Unicycler (v0.4.8) or SPAdes (version 3.13.0) (50). SPAdes includes a step for read error correction and quality trimming of Illumina reads, adding an extra layer of quality control. Various kmer lengths including 21, 33, 55, 77, and 127 were explored along with the ’careful’ mode during SPAdes assembly to ensure high quality assembly. The final genome assemblies were subjected to quality verification through QUAST (47) software to ensure reliable and accurate assembly. This tool integrates confidence scores to enhance the accuracy of the classification.

### Phage clustering and taxonomic analysis

Full phage genomes were clustered using Vcontact2 (51), capable of replicating the genus-level viral taxonomy assignments from the International Committee on Taxonomy of Viruses (ICTV) with 96% accuracy. Blastn (52) was employed to investigate intergenomic similarities within each viral cluster (VC), further delineating relationships between the isolated phages. Phylogenetic analysis was conducted for each VC, constructing neighbor-joining trees based on intergenomic similarity matrices to illustrate the genetic relationships between the phages.

### Phage characterization and genomic organization

The lifestyle of the phages was predicted using the Bacphlip tool (53), which searches for the presence of protein domains associated with the temperate lifestyle in the phage genome. The process of gene calling was carried out through the consensus call of multiple programs, namely Phanotate, Prodigal, Glimmer, and Genemarks, via the integration tool multiPhATE (54). The tRNA genes were specifically identified using tRNAscan (55). Default parameters for gene calling allowed three potential start codons (ATG, GTG, TTG) and three termination codons (TAA, TAG, TGA). A minimum open reading frame (ORF) length was set to 90 nucleotides. The identified genes were annotated with the Phrogs (56) and PDB (57) databases using Hidden Markov Model (HMM) profiles.

### Comparative genomics of the *Klebsiella* phages

Comparative genomics of the isolated phages was done. The genome plot comparison was constructed with the R package gggenomes (58). The pairwise comparisons of all the nucleotide sequences were conducted using the Genome-BLAST Distance Phylogeny method (59) under settings recommended for prokaryotic viruses. The resulting intergenomic distances were used to infer a balanced minimum evolution tree via FastME (60) including subtree pruning and regrafting post processing. Branch support was inferred from 100 pseudo-bootstrap (SPB) replicates each. Trees were rooted at the midpoint and visualized with FigTree (61). Taxon boundaries at the species, genus and family level were estimated with the OPTSIL program (62), the recommended clustering thresholds and an F value (fraction of links required for cluster fusion) of 0.5. The phylogenetic tree of each VC was computed from the similarity matrix using the Neighbor joining algorithm. The resulting trees were represented using iTOL (63). Multiple sequence alignments were constructed using Clustal Omega (64).

### Genetic analysis of phage host recognition

Special attention was dedicated to the genes involved in host recognition, particularly those encoding tail fiber proteins. A RBP detection tool (65) was employed to scan proteins larger than 200 amino acids in size. This method integrates embeddings representations computed through the Prot Trans language model with the identification of the presence of domains associated with tail fiber proteins. This information was then processed by an XGBoost model to predict whether the amino acid sequence indeed corresponds to a tail fiber protein. In addition, the study considered the Dpo domain of proteins, which are usually referred to as tail spike proteins. Due to the sequence divergence yet conserved structures of these proteins (namely β-helix, 6-bladed β-propeller) (33,66), structural investigation was necessary. To facilitate this, proteins whose annotation fell into either tail proteins or unknown proteins were scanned against a local database of HMM profiles derived from Interproscan 5 (67) entries related to EC numbers 4.2.2. and 3.2.1, indicative of carbohydrate lysis activity. Proteins that presented a significant match over a minimum of 50 amino acids were selected, and their 3D structural conformation was predicted using the AI model Esmfoldv1 (68). The presence of the Dpo domain was subsequently confirmed by scanning the predicted 3D structure of the protein against a local Dpo database using foldseek (69). The exact sequence of the Dpo domain was inferred after predicting the domain boundaries using SWORD2 (70).

### Phage host tropism and infectivity

The *Klebsiella* phages were assayed by spot test against the *Klebsiella* spp. strains in order to evaluate their host range in triplicate. Drops of 1 µL of each phage with a titer after its conservation at -70°C >10^5^ PFU/mL were added to bacterial lawns using the soft agar technique (71). Plaques were incubated at 37°C overnight. Strains were considered susceptible to a phage if showed a clear or turbid spot for at least two of the three replicates. In addition, to evaluate infectivity and phage production, spot tests using serial dilutions were performed, using 1:10 and 1:10^3^ dilutions of each phage. In this experiment, single plaques in at least one of the dilutions were considered positive.

### Analysis of the crossed-infection matrix

The obtained crossed-infection matrix was a bipartite matrix, because it contains two types of elements (phages and bacterial hosts) that interact. It can be decomposed into disjoint elements, composed by phages and bacterial hosts that have cross-infections between them but not with other elements (72). Disjoint components of the crossed-infection matrix were identified and plotted using the R package ‘igraph’ (73). Modularity of the matrix was analyzed and the modules were plotted with the R package ‘bipartite’ (74). The statistical significance of the modularity was tested utilizing null models obtained with ‘bipartite’. Six hundred random matrices were used to calculate the significance of our obtained modularity.

### Phage cocktail design and in vitro validation

A phage cocktail was created combining broad-spectrum and narrow-range phages covering the KL-type of at least 45 reference strains, including KLs of some high-risk clones frequently identified in Spanish hospitals causing infections (13). A combination of 12 phages of 10^7^ PFU/mL each was used as a phage cocktail. After its conservation at -70°C, the cocktail was tested *in vitro* by spot tests against the *Klebsiella* spp. reference strains as previously described and against a panel of 58 carbapenem-resistant *K. pneumoniae* clinical isolates. In addition, analyses of the association between susceptibility to the phage cocktail and other qualitative variables in clinical isolates were performed. The correlation between different qualitative variables of the clinical isolates (susceptibility to the cocktail, KL, ST, O-Ag and carbapenemase-producing genes) was studied with a Pearson’s χ^2^ analysis or Fisher’s exact test depending on expected frequencies using R commander package (71).

### Anti-phage defense analysis of clinical isolates

Although phage recognition and phage adsorption are key for the infectivity, post-entry mechanisms are also important for successful phage replication and bacterial lysis. To evaluate the possible correlation of anti-phage defense systems with resistance to the phage cocktail, anti-phage defense mechanisms were identified using defense-finder (39) from the sequences of the clinical isolates used in this study. CRISPR-Cas systems were detected using CRISPRCasTyper (75). Spacers were compared with phages from the cocktail using blastn (52). Differential probability of resistance (dPR) was calculated as de difference between the number of isolates with a defense mechanism (DF) that are resistant to the cocktail (R) and the number of isolates with this DF that are susceptible to the cocktail (S): dPR = P(R|DF) - P(S|DF). To avoid the effect of the association of some defense-systems with a particular KL, this was calculated separately for clinical isolates with potentially susceptible KL, in which at least one clinical isolate with this particular KL was susceptible to the cocktail but others were resistant. Values close to 1 were considered as possibly related with resistance to the cocktail.

## Results

### Phage diversity and distribution in viral clusters

Extensive phage hunting from environmental samples in Valencia (Spain) was done using the 77 *Klebsiella spp*. reference strains as primary hosts. A total of 83 phages were isolated, and among them, 71 were successfully sequenced and characterized. Each of the strains was infected by at least one of the phages, suggesting the possibility to isolate *Klebsiella* spp. phages from the environment capable of infecting a large collection of *K. pneumoniae* isolates with different KL types. Three phages of the genus *Sugarlandvirus* previously isolated in our group were included in the phage collection (76). The genome length of the phages varied significantly, ranging from 38,774 to 176,687 base pairs (Supplementary material 2). The isolated phages were distributed across 19 distinct viral clusters (VC) corresponding to several genera, with a confidence level higher than 0.9 (Figure 1). These included *Webervirus* (18 phages), *Przondovirus* (15 phages), *Drulisvirus* (9 phages), *Vectreviru*s (5 phages), *Mydovirus* (4 phages), *Sugarlandvirus* (3 phages (76)), VC-818-0 (3 phages), VC-473-0 (1 phage), VC-590-0 (2 phages), VC-589-2 (1 phage), and *Efbeekayvirus*, *Teetrevirus*, *Slopekvirus*, *Aerosvirus*, *Taipeivirus*, *Jiaodavirus*, and *Nonagvirus* (each with a single phage). Four remaining phages were categorized as outliers.

**Figure 1.**
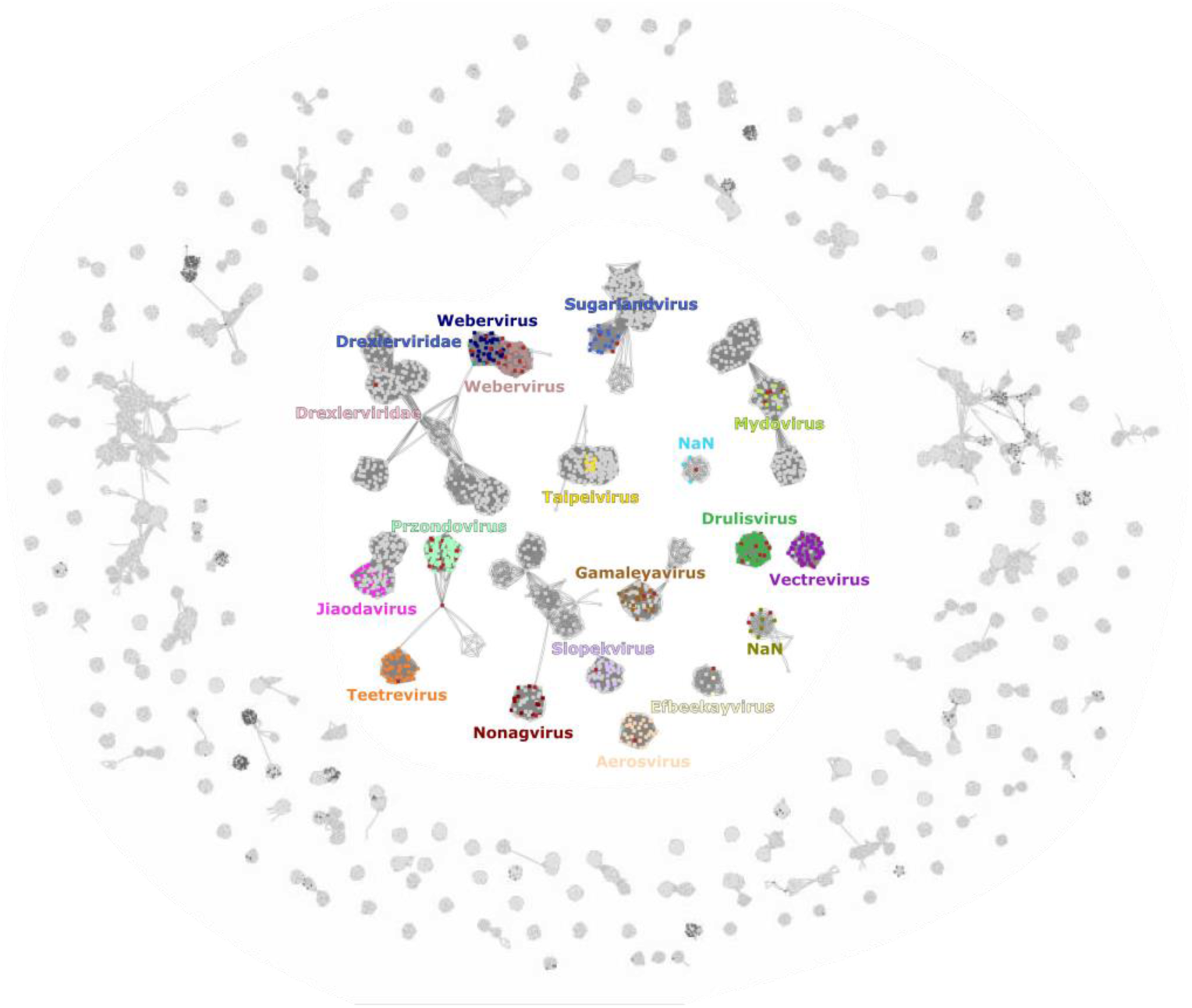
Viral clusters network including 76 newly isolated and characterized *Klebsiella* phages. The network analysis has been performed on protein profiles encoded by phages using the vConTACT2 software. Each of our *Klebsiella* phage isolates are represented as red nodes. The phages that belong to the same viral cluster as our phages were represented with specific color nodes. The gray nodes correspond to other phages that were used as references for our analysis. The graph was visualized using a force-directed layout through the Cytoscape software (83). Nan: non available name.

### Phage genomic characterization and annotation

The lifestyle and the virulence of the phages was predicted obtaining a probability value ranging from 0 to 1, which indicates its probability to exhibit a virulent phenotype. All isolated phages were predicted as lytic life cycles, with the probability ranging from 0.72 to 1 for over half of the phages, being the mean lytic life predicted probability for an isolated phage 0.96 ± 0.05. Although genetic annotations were carried out using the remote homology search tool, HMMer, in conjunction with Phrogs database (56,57,77), a large fraction of genes remained with an unknown function (Supplementary material 3). Core genes across the phage genomes were categorized into defined functional groups. These included the large and small terminase subunits and the major head protein in the ’Head and Packaging’ group, HNH endonuclease and exonuclease in the ’DNA, RNA and Nucleotide Metabolism’ category, the spanin, holin, and endolysin in the ’Lysis’ category, and the prototypical tail proteins and tail length tape measure protein in ’Tail Proteins’, along with tail fiber and tail spike proteins in the ’Receptor Binding Proteins’ category (Figure 2). Alongside these conventional genes, our analysis also highlighted less conventional genes, like transcription regulation genes, and others associated with overcoming bacterial defense systems.

**Figure 2.**
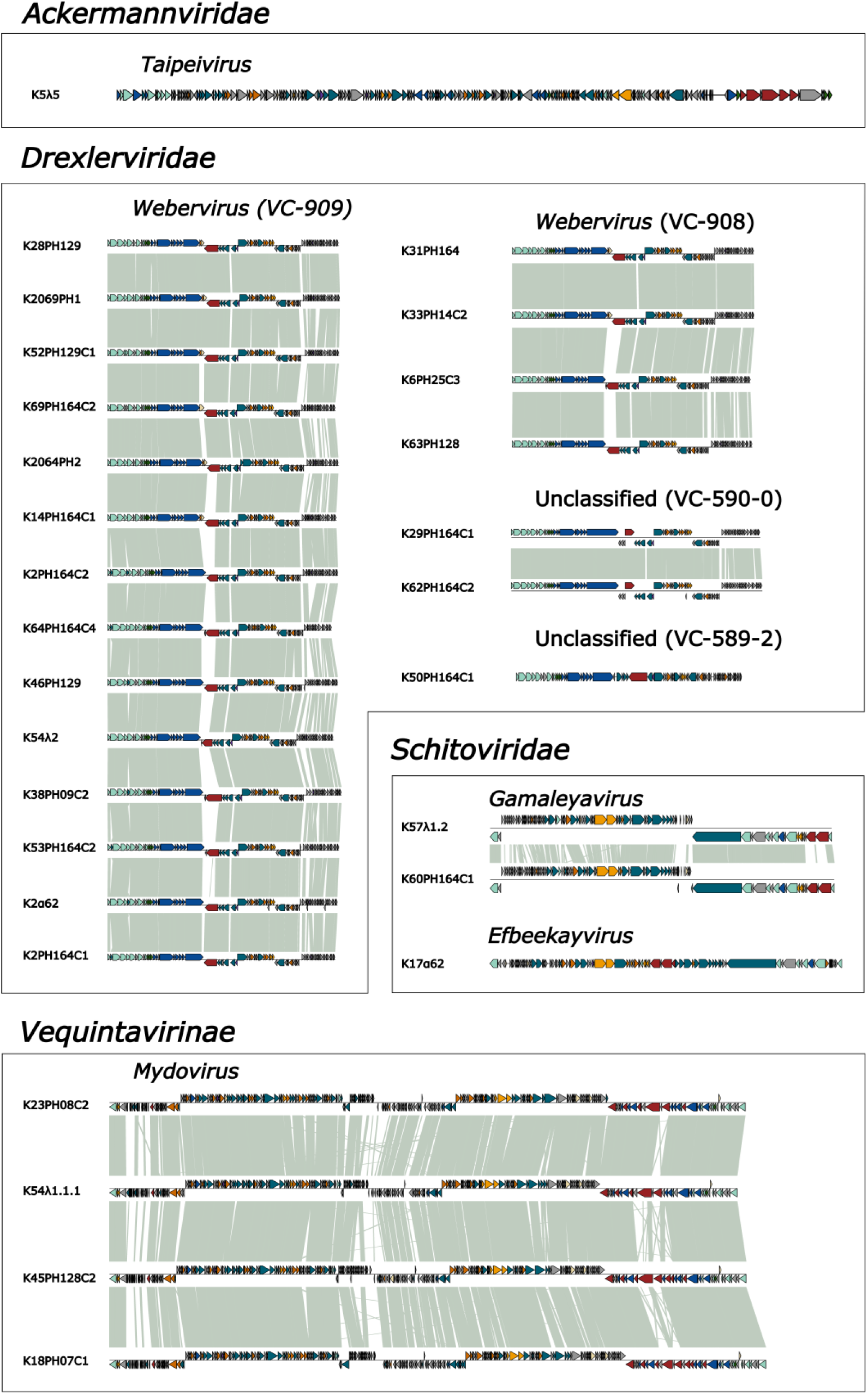
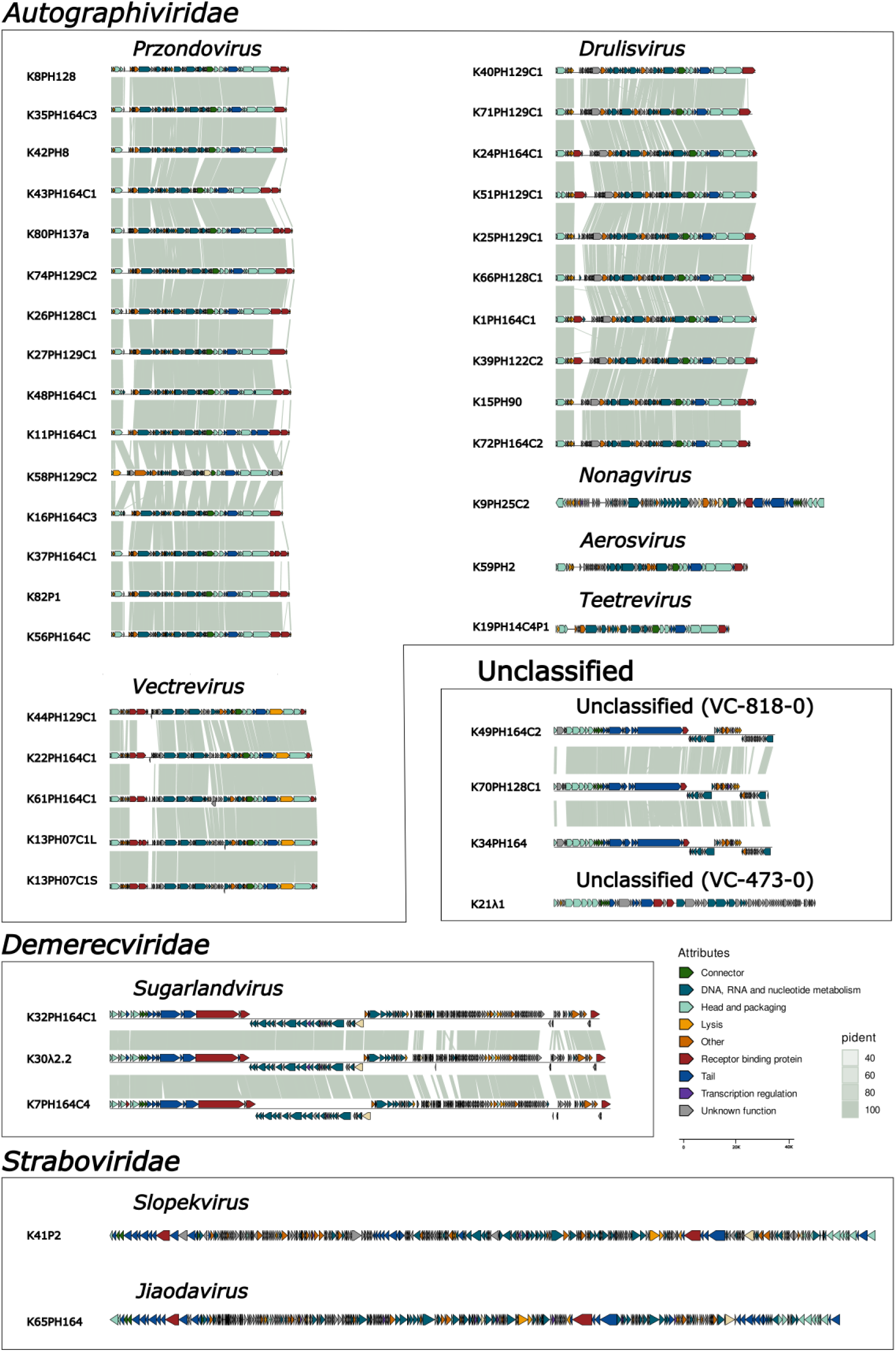
Intergenomic similarity between *Klebsiella* phages taxonomically classified. The VCs obtained after the vContact2 process were organized in family or sub-family and genus. The intergenomic similarities were computed using blastn (52). The arrows, oriented along the coding strand, represent the identified opening reading frames. Each color is related to a functional group, as defined in the Phrogs database (56). Correspondence between the name of the phages and the alias is available in Supplementary Material 1.

**Figure 3.**
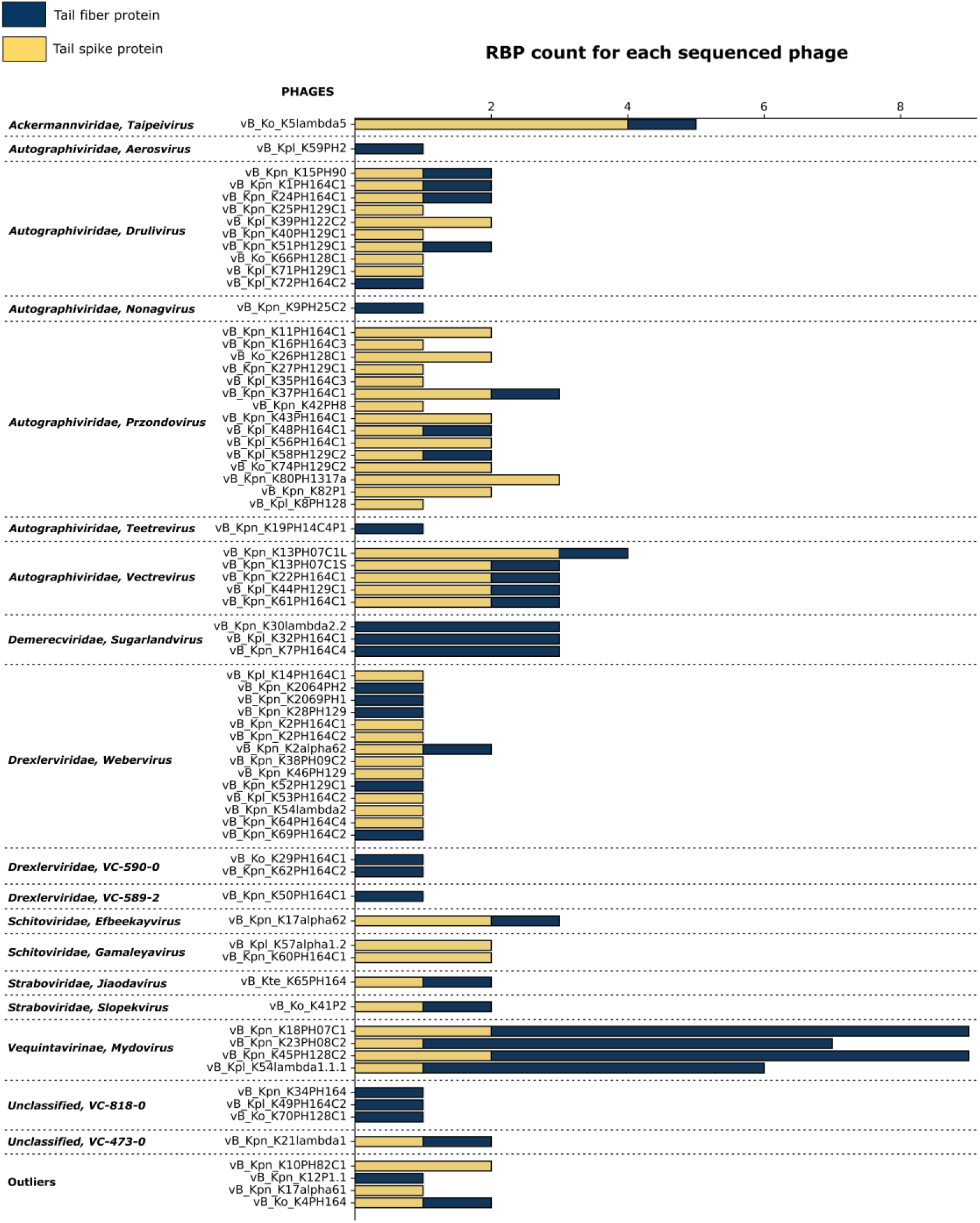
Count of the RBPs in each isolated *Klebsiella* phages arranged by viral cluster. The chart plot represents in the vertical axis each of the phages classified in their viral clusters. The horizontal axis indicates the total RBP count. The count of tail fibers is represented in blue, and the count of tail spikes in gold.

### Phage host recognition genes

In order to determine phage host recognition genes, a total of 150 RBPs were identified, which included 79 tail spike proteins. The diversity and distribution of these RBPs, however, varied significantly across different VCs (Supplementary material 4). Every isolated phage contained at least one identifiable RBP. Within most VCs, we observed a mixture of tail fiber proteins and tail spikes among the identified RBPs. The highest count of tail spikes was four, in phage vB_Ko_K5lambda5. However, certain VCs, such as VC-184-0, VC-590-0, and VC-818-0, exhibited no identifiable tail spike proteins. The structural variation in the identified Dpo domains was remarkable. The right-handed β-helix was the most prevalent fold, representing 75 of the identified Dpo domains, followed by the 6-bladed β-propeller with 4 domains. The structural architecture of the tail spike proteins was also diverse and organized into one or more domains. The simplest form was a single domain Dpo with either a β-helix or a 6-bladed β-propeller, whilst other were arranged into two domains. More complex architectures included three domains, consisting of an N-terminal domain, a right-handed β-helix, and a jelly roll. Some tail spikes displayed even more intricate structures, like multiple domain organization or baseplate assemblies with a 6-bladed β-propeller. Interestingly, structural diversity was also observed within the right-handed β-helix of Dpo domains, exhibiting variabilities in length. Furthermore, our analyses suggested a structural continuity between some tail fiber proteins and tail spike proteins. In fact, instances of tail fiber proteins that comprised the phage T7 N-terminal domain, an α-helix and the jelly-roll motif were observed. Both domains demonstrated structural similarity when aligned with tail spike proteins (Supplementary material 5).

### Determination of phage tropism and infectivity

Host range of the 86 isolated phages was determined experimentally by a crossed-infection matrix including the 77 *Klebsiella* spp. reference serotypes strain collection (Figure 4A). The obtained crossed-infection matrix consisted of 6,622 interactions, being only 3.47% of them (230/6,622) positive. In general, most phages were highly specific, being able to infect a small amount of hosts (average of 2.67). In addition, hosts tend to be better resistant than permissive, being susceptible to few phages (average of 2.99) (Figure 4B). Analyzing the modularity (Q) of the matrix, we identified 17 separate modules. The overall modularity of the matrix (Q=0.604) was slightly higher than the average modularity observed in 600 realizations of the null model (Q=0.572), and similar to the highest of them (Q=0.604). Most positive interactions were found within the modules (180/230, 78.26%) (Figure 4A). The whole matrix was composed of a total of 17 disjoint elements, sets of phages and hosts with cross infections within them but not with other elements (Supplementary material 6). The elements were not related to viral families or other taxonomic relationships.

**Figure 4.**
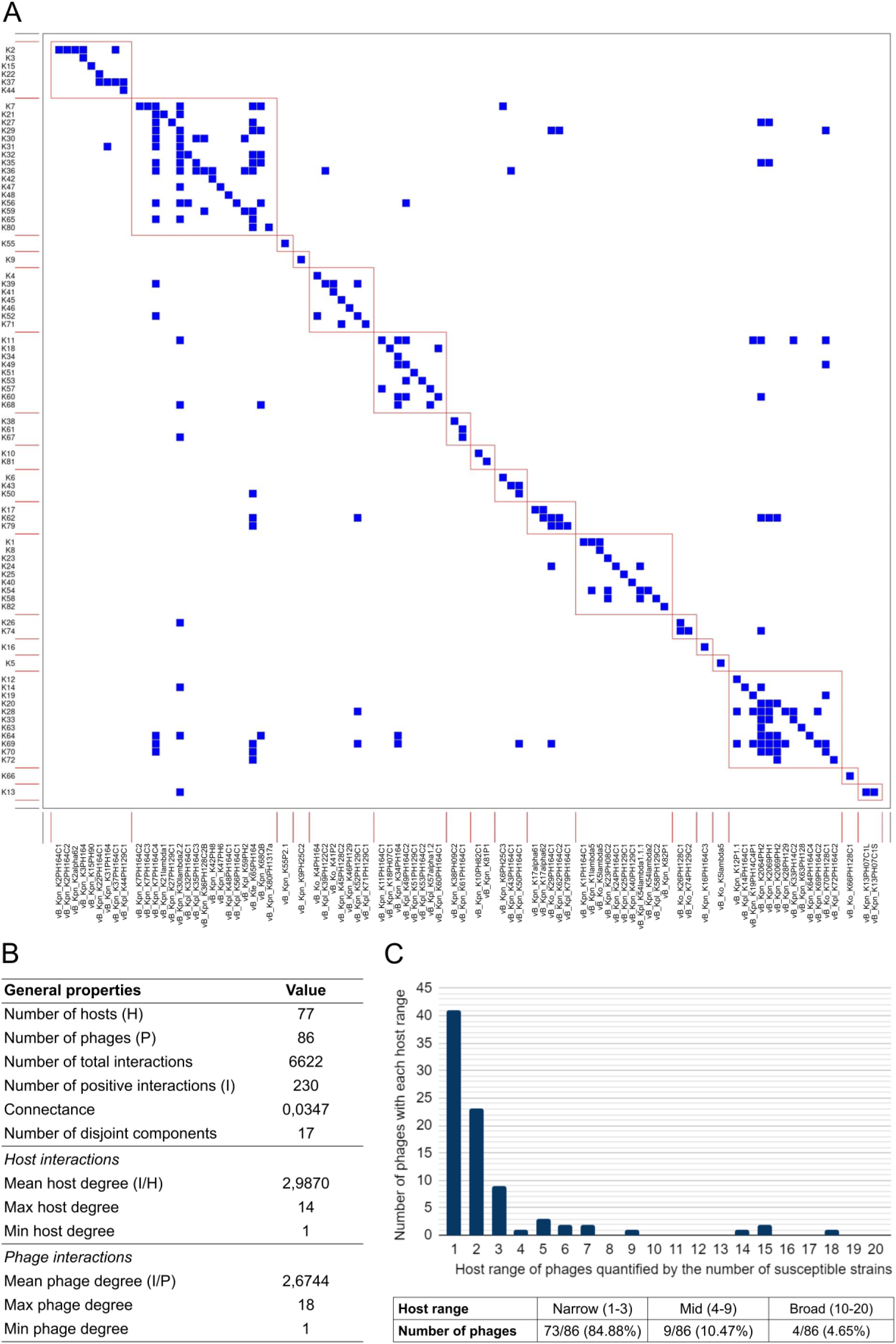
Host range of 86 *Klebsiella* phages against 77 *Klebsiella* reference serotypes. (A) Modular distribution of the crossed-infection matrix representing the host range of the 86 *Klebsiella* phages in each row over the 77 *Klebsiella* spp. reference strains in each column. Positive interactions are represented in blue and negative interactions are represented in white. 17 modules are detected and squared in red. Modularity value (Q) is 0.604 (p-value=2.2×10^-16^). (B) General properties of the phage-bacteria crossed-infection matrix host interactions and phage interactions. (C) Graphic representation of the diverse host range of the 86 *Klebsiella* phages. The horizontal axis represents phage host range quantified by the number of susceptible reference strains, while the vertical axis represents the amount of *Klebsiella* phages infecting each quantity of *Klebsiella* spp. reference strains. We consider narrow host range for phages that infect less than 4 strains, mid host range for phages that infect between 4 and 9 strains, and broad host range for phages that infect up to 10 strains. Rate and percentage of phages in each category are available in the table below the graph.

Focusing on the diversity of phage host ranges in our phage collection infecting the reference strains, 73 of the 86 isolated phages (84.88%) showed narrow host range, being able to infect between 1 and 3 strains. Nine phages (10.47%) were mid host range, being able to infect between 4 and 9 strains. Only 4 phages (4.85%) exhibited broad host range, being able to infect more than 8 strains (Figure 4C). In most VC, the HR was consistent across the phages with a standard deviation (std) lower to 1 besides the VC-184-0 (std = 6.94) and the VC-909-0 (std = 3.77).

### Phage cocktail design

Phage cocktails can be an interesting approach to implement ready-to-use preparations for prevention or to treat acute infections. Under this scenario, we designed a cocktail encompassing a combination of 12 phages, as suggested by the Regulatory Medicine and Sanitary Products Agency of Spain (AEMPS) as the maximum recommended number of phages for a single cocktail. Based on the data obtained for the phage collection reported here, we included vB_Ko_K5lambda5, vB_Kpn_K7PH164C4, vB_Kpl_K8PH128, <colcnt=3> vB_Kpn_K50PH164C1, vB_Kpn_K30lambda2.2, vB_Kpn_K34PH164, vB_Kpn_K45PH128C2, vB_Kpl_K54lambda1.1.1, vB_Kpl_K44PH129C1, vB_Kpn_K60PH164C1, vB_Kte_K65PH164 and vB_Ko_K74PH129C2. Firstly, the phage cocktail was tested *in vitro* by serial dilutions of the cocktail using the spot test method in semi-solid media over each of the 77 *Klebsiella* spp. reference strains (Supplementary materials 7A). Forty-two of them showed susceptibility to the cocktail (54.55%), as shown by the single plaques observed in the dilutions tested. Expected host range of the cocktail considering the sum of single phages host range was 45 strains (58.44%) (Supplementary materials 7B). Discrepancies between the expected and observed results were observed for five of the strains. Indeed, *Klebsiella* spp. reference strains of the capsular serotypes 14, 21, 26 and 47 (K14, K21, K26 and K47) were not able to be infected, although it was expected to observe plaques even at low efficiency of plating. In contrast, strain with capsular serotype 41 (K41) was not expected to be infected, while single plaques were observed during the experiments (Supplementary materials 7A).

### Carbapenem-resistant clinical isolates representing *K. pneumoniae* high-risk clones

A collection of 58 carbapenem-resistant *K. pneumoniae* high-risk clones circulating in Spain was used. The genomic analysis showed that the clinical isolates belonged to 10 KL-types, including six reference KLs (KL25, KL17, KL24, KL27, KL64, KL48), and 4 non-reference KLs (KL102, KL107, KL112, KL151), and thus not included as primary hosts during our phage hunting approach. The panel represented 8 STs, 5 O-locus, and 4 carbapenemase types (OXA-48-like, KPC, VIM, NDM) (Supplementary material 6). The KL-type was significantly associated with the ST of each strain (χ^2^=315.19; p<0.0001), as previously known due to the clonal expansion of *K. pneumoniae* isolates. Indeed, 6 of the 8 ST represented in our panel were associated with a single KL-type. Only ST15 and ST11 were associated with four and five KL-types respectively (Figure 5A). In addition, a significant association between those parameters and the O-locus (OL) was detected (ST-OL: χ^2^=114.81; p<0.0001, KL-OL: χ^2^=171.55; p<0.0001). All isolates with ST101-KL17, ST147-KL64, ST11-KL64, ST11-KL24 and ST15 in our panel had OL-type O1/O2v1. Also, all isolates with ST307-KL102 and ST512-KL107 had OL-type O1/O2v2, and all isolates with ST392-KL27 and ST405-KL151 had OL-type O4. The only isolate with OL-type O5 was ST11-KL25, and the only one with OL102 was ST11-KL107. Regarding carbapenemase-encoding genes, *bla*_OXA_ and *bla*_KPC_ were significantly associated with ST, KL and OL (*bla*_OXA_-ST: Fisher’s exact test p<0.0001; *bla*_OXA_-KL: Fisher’s exact test p=0.0005; *bla*_OXA_-OL: Fisher’s exact test p<0.0001; *bla*_KPC_-ST: Fisher’s exact test p<0.0001; *bla*_KPC_-KL: Fisher’s exact test p<0.0001; *bla*_KPC_-OL: Fisher’s exact test p=0.0072), but it did not occur for *bla*_NDM_ and *bla*_VIM-1_ (*bla*_NDM_-ST: Fisher’s exact test p=0.6257; *bla*_NDM_-KL: Fisher’s exact test p=0.2537; *bla*_NDM_-OL: Fisher’s exact test=0.1047; *bla*_VIM-1_-ST: Fisher’s exact test p=0.3646; *bla*_VIM-1_-KL: Fisher’s exact test p=0.5690; *bla*_VIM-1_-OL: Fisher’s exact test p=0.1708). All isolates from our panel with ST101-KL17-O1/O2v1 were positive for *bla*_KPC-2_, and all with KL107 for *bla*_KPC-3_ except KL107-ST512-O1/O2v2, which was positive for *bla*_KPC-23_. In contrast, any isolates with ST15, ST405-KL151-O4 or ST11-KL25-O5 were positive for any *bla*_KPC_ gene. In addition, all isolates with ST405-KL151-O4 and ST392-KL27-O4 were positive for *bla*_OXA-48_, and any with ST101-KL107-O1/O2v1, KL17 or ST11-KL25-O5 were positive for any *bla*_OXA_ gene (Supplementary materials 1). In addition, anti-phage defense systems profile for each clinical isolate was also evaluated, finding 49 different subtypes. Each isolate presented between 7 and 21 different subtypes of defense systems. CRISPRs were detected in 27 isolates, but spacers did not align with any of the phages present in the cocktail (Supplementary materials 8).

**Figure 5.**
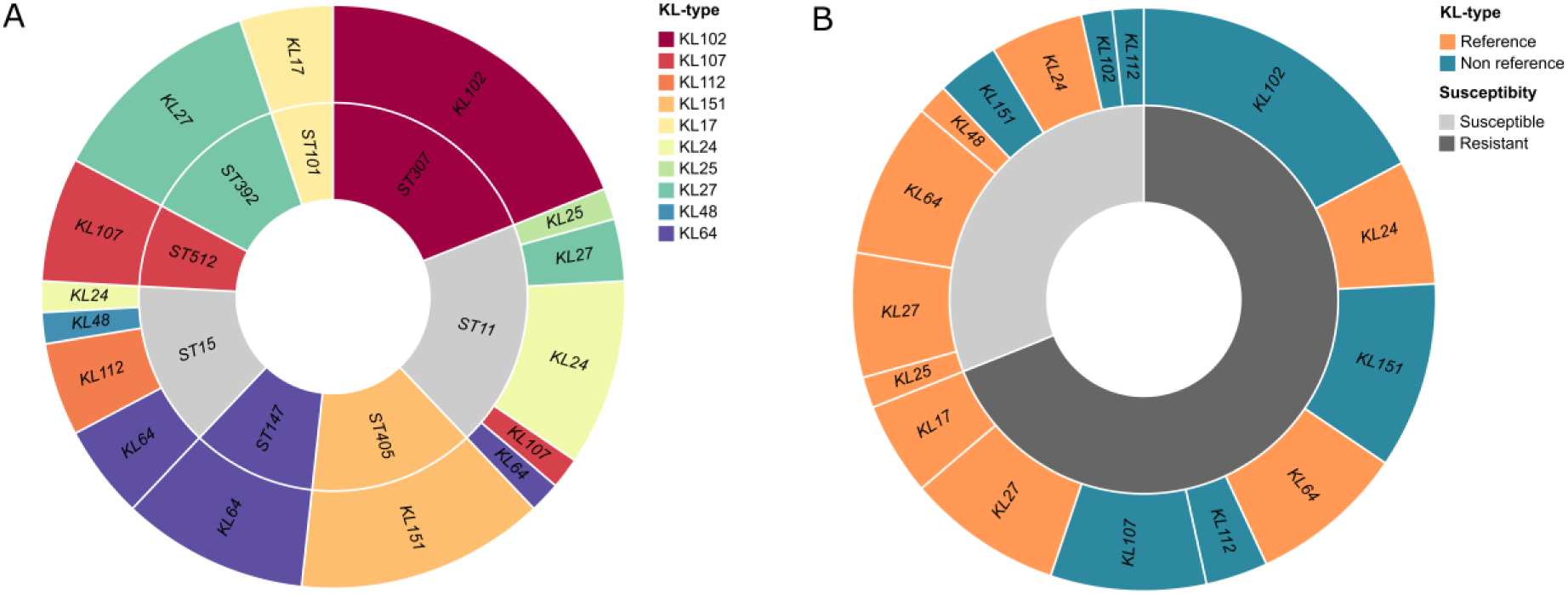
Association between KL, ST and susceptibility to the phage cocktail in a panel of 58 carbapenem-resistant *K. pneumoniae* clinical isolates. (A) Association between KL and ST. In the inner circle are represented the different ST of the isolates proportionally to the number of isolates with each ST. In the outer circle there are represented the different KL of the isolates with each ST, also proportionally to the number of isolates with each KL. Colors represent the different KL. ST colored in gray are not associated with a single KL. (B) Association between KL and susceptibility to the phage cocktail. The inner circle represents the proportion of susceptible and resistant isolates, represented in light gray and dark gray respectively. In the outer circle are represented the different KL of the strains susceptible and resistant to the cocktail. Some KL are present in both categories because there are both resistant and susceptible isolates with these particular KL. Portions are proportional to the number of isolates resistant or susceptible with each particular KL. Reference KL-types are represented in orange, and non-reference in blue. 77.8% of susceptible isolates have a reference KL-type, while only 22.2% have a non-reference KL-type. In contrast, 42.5% of resistant isolates had a reference KL-type, while 57.5% had a non-reference KL-type. This graph was made using RawGraphs (84).

### Phage cocktail validation in high-risk clones

To study the implications of a phage cocktail in the biocontrol of *K. pneumoniae*, the designed cocktail was assessed *in vitro* against the panel of 58 carbapenem-resistant clinical isolates. The results showed that the cocktail was able to infect 18 of them (31%) (Supplementary materials 1). To further understand these results, we evaluated the KL-types. Only 31 of the 58 had a KL-type included in the 77 *Klebsiella* spp. reference serotypes collection (reference KL-type), and 27 had a KL-type not included. Going further, only 14 of the 31 isolates with reference KL-types were infected by the cocktail (45.16%), as well as 4 of the 27 isolates with non-reference KL-types (14.81%) (Figure 5B). The results showed that the relationship between having a reference KL-type and being infected by the cocktail was significant (χ^2^=7.99; p=0.0127). Considering exclusively the 31 isolates with reference KL-types, it is worth mentioning that KPN04 (KL25) and KPN19 (KL48) were susceptible to the cocktail, despite reference strains with KL-types KL25 and KL48 were resistant to the cocktail. The three KL17 isolates were also resistant to the cocktail, as well as KL17 reference strain. In addition, 3/7 KL24 (42.9%), 4/9 KL27 (44.4%) and 5/10 KL64 isolates (50%) were susceptible, as the KL24, KL27 and KL64 reference strains. Regarding non reference KL-types, all the KL107 clinical isolates were resistant to the cocktail, and only 1/11 KL102 (9.1%), 1/3 KL112 (33.3%) and 2/8 KL151 (25%) were susceptible (Supplementary materials 9).

Despite having a KL-type included in the reference strains collection was associated with susceptibility to the cocktail, the KL itself did not seem to affect significantly (Fisher’s exact test p=0.1181), as well as the ST (p=0.1028). OL was neither significantly related with susceptibility (Fisher’s exact test p=0.2390), as well as encoding *bla*_OXA-48-like_, *bla*_NDM_, *bla*_KPC_ or *bla*_VIM_ carbapenemase genes (Fisher’s exact test p=0.7027; Fisher’s exact test p=1; Fisher’s exact test p=0.3473; Fisher’s exact test p=0.055) (Supplementary materials 9).

The role of anti-phage defense systems in resistance to the cocktail was thus explored. Some of those defense systems may be related with an increased resistance to the phage cocktail, however, data were not conclusive enough to establish a significant correlation between resistance to the cocktail and any of the detected defense systems (Supplementary material 10).

### Targeted phage hunting in clinical isolates and phage characterization

Given the poor infectivity of our designed phage cocktail against clinical isolates, we decided to explore if phage hunting directed to specific clinical isolates would be a better approach to combat *K. pneumoniae* nosocomial infections. To achieve this goal, we selected KL64 clinical isolates, since this KL type was highly represented in our panel of clinical isolates (10 isolates), but only half of them were susceptible to our phage cocktail. Our bioprospecting allowed us to isolate 20 new phages able to infect KL64 isolates. Seventeen of them were successfully sequenced, phylogenetically classified and functionally annotated (Supplementary materials 11). A crossed-infection matrix was performed by spot test including these new 20 phages and phage vB_Kpn_K64PH164 (isolated in the K64 reference strain), and the 11 KL64 bacterial hosts (Figure 6). A total of 231 interactions were tested in triplicate, being 48% of them positive (112/231), which is remarkable given the 3.48% of positive interactions in the previous matrix. Phages were able to infect between 1/11 to 10/11 (average of 5.33) different isolates. Hosts were susceptible to an average of 10.18 phages. The most resistant isolate was susceptible to 4/21 phages, and the most permissive was KL64 reference strain, susceptible to all phages (21/21).

**Figure 6.**
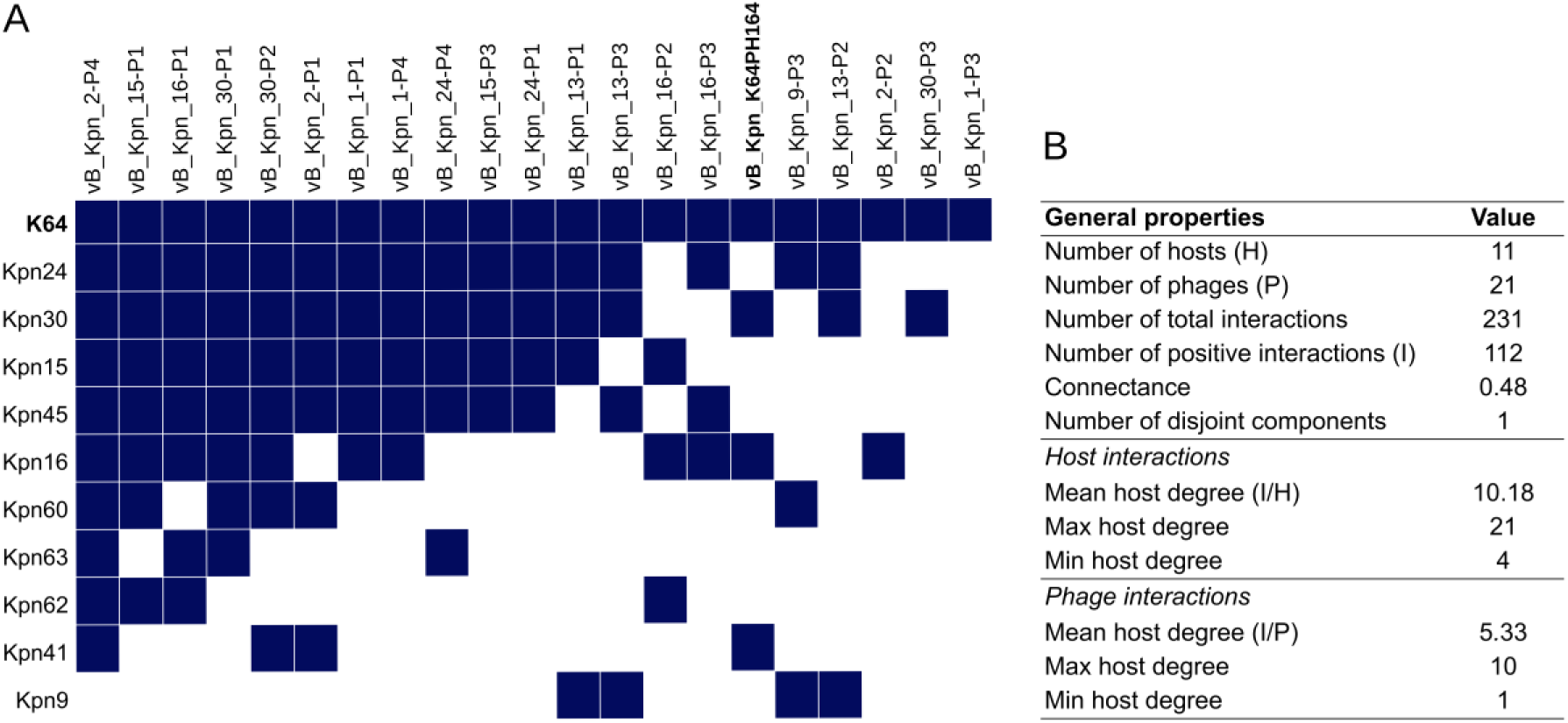
Host range of 21 *Klebsiella* phages and 11 isolates with K-type 64. (A) Crossed-infection matrix between 21 *Klebsiella* phages and 10 clinical isolates with K-type 64 and the K64 reference strain. 20 of the phages have been isolated in KL64 clinical isolates and one in the reference strain K64. K64 and phage isolated in this reference strain are represented in bold. (B) General properties of the phage-bacteria crossed-infection matrix host interactions and phage interactions.

## Discussion

In this study, we have implemented a systematic phage hunting that allowed us to successfully isolate a large collection of new *Klebsiella* phages, showing an invaluable diversity. Previous research has indeed documented the construction of *Klebsiella* phage banks and examined the host range of phages across bacterial collections (57,78,79). However, our collection distinguishes itself as the first to include genomically characterized phages with the potential to infect each of the 77 *Klebsiella* spp. reference strains, thus encompassing all known *K. pneumoniae* capsular serotypes. Our findings revealed an extensive genomic diversity among the isolated phages, as they were categorized into more than 19 distinct VCs, in addition to four outlier phages representing unknown variability that may be representatives of potential new genera. This heightened diversity could be attributed not only to the high number of isolated phages, but also to the isolation strategy. By employing a panel of bacteria highly diverse in capsular serotypes, a crucial determinant of phage infection, a more expansive diversity in the isolated phages was unveiled. Our *Klebsiella* phage bank is one of the widest and diverse collections of genomically characterized phages with known host ranges tested in a capsule diverse strain collection. Thus, it is a valuable resource for personalized phage therapy, and a useful dataset for the implementation of prediction tools for phage-host interactions using machine learning approaches.

A crucial step in the infection process involves phage recognition to its host, a process largely mediated by RBPs, including tail fibers and tail spikes (21,80). By using the *Klebsiella* spp. reference strains with different capsular types for phage isolation, we have been able to capture significant diversity in tail spikes. The presence or absence of tail spikes appeared to be a shared feature among phages from the same viral cluster. However, some exceptions were found in the *Webervirus,* where a Dpo domain was undetectable in some phages while the majority carried at least one. Nonetheless, a halo could be observed around the lytic plaques formed by those phages, indicating potential Dpo activity (81). This suggests that the current methods may not have been sensitive enough to identify Dpo, warranting improved methods for their detection (26). While the host range of the isolated phages was diverse, it appeared to be conserved within a viral cluster, besides our isolated phages of genera *Webervirus* and *Sugarlandvirus.* Most of the broad host range phages carried one or no Dpo domain, suggesting the possibility of an infection mechanism that is not dependent on the nature of the capsule (26,76)

The high specificity of most *Klebsiella* phages challenges the implementation of phage-based broad-spectrum preparations to fight and prevent infections caused by MDR *K. pneumoniae*. In this work, we explored the possibility of designing a broad-spectrum cocktail able to infect a wide range of *Klebsiella* KL-types. The number of phages included in a phage cocktail should be determined by the desired host range of the preparation, and also by possible phage-phage interactions (43). The limited number of broad-range phages in our collection forced us to design a phage cocktail to cover the *Klebsiella* spp. reference strain collection. To evaluate the biomedical implications of our phage cocktail, we tested it in a collection of clinical isolates of carbapenem-resistant *K. pneumoniae* circulating in Spanish hospitals. Our panel included representatives for some of the most problematic high-risk clones (ST307, ST11, ST512, ST15, ST147 and ST392), some of them reported worldwide (13). The strong correlation observed of most ST with a single KL-type has been also reported in a recent study about the European distribution of carbapenemase-producing *K. pneumoniae* (12). Indeed, ST512 was associated with KL107, and ST101 with KL17, as observed in our panel. This correlation between KL and ST may be due to the impact of local clonal expansions of this pathogen (12). Here, only 31% of clinical isolates were susceptible to the cocktail, and the variable associated with susceptibility was sharing or not a reference KL-type, having 45.2% of infection considering exclusively reference KL-type isolates. Indeed, phages from the cocktail were isolated using *Klebsiella* spp. reference strains, so the association correlated with previous works which consider *Klebsiella* KL-type as the main determinant for phage tropism (28). However, our results revealed that even if a reference capsular type is infected by the cocktail, not every single isolate sharing KL-type will be infected. This suggests that post-entry mechanisms may contribute to the resistance to the phage cocktail (28,39,40,82). In addition, our results evidence the limitations of elaborating broad-range cocktails based on phages isolated from reference strains. Clinical isolates seem to be much more restrictive to phage infection than the reference strains, probably due to their acquisition of more complex and diverse defense systems that avoid the infection of a high proportion of those phages able to target its KL-type. In this sense, we explored the possibility to isolate phages directly through the clinical isolates. We suggest that phage hunting directed to specific *K.* pneumoniae clinical isolates is an efficient antimicrobial control strategy, and may be the best approach to implement personalized anti-*K. pneumoniae* treatments.

We are aware that our study presents some drawbacks. Preparation of ready-to-use phage cocktails might be an interesting strategy to reduce the bacterial burden, as preventive tools. However, reference strains have been suggested to be used as primary hosts for phage hunting and phage production, due to the lower levels of toxins and others. A major limitation in *Klebsiella* spp. is that the reference capsular types collection is based on serotyping, which is limited to a few KL-types, excluding a large amount of variability and reducing the possibilities to isolate therapeutic phages. In addition, clinical isolates have acquired post-entry defense systems that can prevent infectivity. However, we can solve this issue isolating phage directly through the clinical bacterium, as suggested here using our systematic approach. Interestingly, and thanks to the permissiveness of the reference strains, we can use them to amplify and produce the phages for therapeutic purposes. As for our collection of clinical isolates, it includes a reduced number of isolates, which may bias our descriptive statistical analyses, but it includes high-risk clones that have been identified as representing the most problematic ST and KL in *K. pneumoni*ae clinical infections (12,13). Another relevant limitation is the evaluation of our experimental results by spot tests, assays presenting qualitative data which sometimes presents discrepancies compared with other techniques like liquid infections (28). Despite the constrains mentioned, this work represents the larger collection of fully characterized *Klebsiella* phages and the first evaluation of a broad-range phage cocktail designed based on the phage-host interactions encompassing all the reference KL-types.

## Conclusions

*K. pneumoniae* is a highly variable encapsulated bacterium, representing a major challenge for designing phage-based products to prevent or treat pathogenic infections. Our results demonstrate the potential to isolate phages for any high-risk clone, suggesting the interest in developing phage hunting directed to specific *Klebsiella* spp. isolates as a faster and more efficient way to implement phage therapeutics as personalized tools. This would allow the design of adapted phage cocktails based on the epidemiology of the region, with a few phages targeting the clones that cause the majority of infections. In addition, phage production and scalability could be achieved using reference strains, which are more permissive and easier to manipulate. Our results are a step forward for new phage-based strategies to control *K. pneumoniae* infections, highlighting the importance of understanding phage-host interactions to design rational cocktails as personalized treatments against *Klebsiella* spp.

## Acknowledgements

We thank Laboratorio de referencia para tipado molecular de patógenos nosocomiales y detección de mecanismos de resistencia a antimicrobianos de interés sanitario de Andalucía (Hospital Virgen Macarena, Sevilla, Spain), Pilar Barberán for help with phage hunting, and Amanda Martínez for technical assistance. This research was funded by ESCMID Research Grant 20200063, project PID2020-112835RA-I00 funded by MCIN/AEI/10.13039/501100011033, and project SEJIGENT/2021/014 funded by Conselleria d’Innovació, Universitats, Ciència i Societat Digital (Generalitat Valenciana) to P.D.-C. C.F.-G. was funded by a PhD fellowship Atracció de Talent UV-INV_PREDOC-1913324 from Universitat de València. R.C.-E. was funded by Grant GRISOLIAP/2020/158 from the Conselleria d’Innovació, Universitats, Ciència i Societat Digital (Generalitat Valenciana) to R.S. S. G.-C. was recipient of a grant from the Research Talent Attraction Program - Modality 1 funded by Comunidad de Madrid (2018-T1/BMD-11174 and 2022-5A/BMD-24243). P.D.-C. was financially supported by a Ramón y Cajal contract RYC2019-028015-I funded by MCIN/AEI/10.13039/501100011033, ESF Invest in your future.

## Author contributions

C.F.-G. performed the experiments, contributed to computational analyses, data analysis, visualization, and manuscript writing. R.C.-E. contributed to computational analyses, visualization, and manuscript writing. M.B.-G. contributed to experiments. F.F.-C. contributed to bacterial resources, computational analyses and manuscript revision. S.G.-C. contributed to bacterial resources, computational analyses, and manuscript revision. J.E.C.-G. contributed to bacterial sequence analyses. R.S. contributed to designing research, and manuscript revision. P.D.-C. provided reagents, conceived the project, designed the research, contributed to data analysis, revised the manuscript, and conducted the supervision. All authors read and approved the final manuscript.

## Declaration of interests

The authors declare no competing interests.

## Supplementary Materials

Supplementary material 1**. Description of the panel of carbapenem-resistant *K. pneumoniae* clinical isolates.** Name and code of the isolates, source, sample type, KL-type, ST, carbapenemase-producing genes and susceptibility to the phage cocktail.

Supplementary material 2**. Isolated and sequenced *Klebsiella* phages.** Genome length, sequencing depth, GC content, number of CDS identified, taxonomy, source, accession number of the isolated and sequenced *Klebsiella* phages and list of susceptible KL types.

Supplementary material 3. **Functional annotation of *Klebsiella* phages.** Protein-coding gene prediction source for each CDS, start and end position of each one, the DNA strand location and the functional annotation of the sequenced *Klebsiella* phages.

Supplementary material 4. **Architectures of the identified tail spike proteins in the isolated *Klebsiella* phages.** 3D representation of the detected architectures of the tail spike proteins in the isolated phages. (A) The single domain tail spikes. (B) The two domains of tail spikes. (C) the three domains of tail spikes. (D) The complex identified tail spikes. In red the right-handed β-helix, in blue the 6-bladed β-propeller, in gold the jelly-roll, in green the N-terminal domain T7.

Supplementary material 5. **Illustration of the structural continuity between the tail fiber and the tail spike proteins.** 3D structure representations and alignments of RBPs identified in the isolated phages. In red: the right-handed beta-helix, in blue: the tail fiber protein; in gold: the tail spike protein. (A) Alignment of two protein units (protein unit 1: 1-76; protein unit 2: 77-250) of the tail fiber vB_Kpn_K18PH07C1, CDS 246 and the tail spike vB_Kpl_K39PH122C2, CDS 55 using the Needleman–Wunsch algorithm under Chimerax (85). (B) 3D representation of the tail fiber vB_Kpn_K18PH07C1, CDS 246.

Supplementary material 6. **Network representation of the crossed infection matrix.** We observe 17 disjoint components. Blue nodes represent hosts and orange nodes represent phages. Numbers in orange nodes correspond to these phages: 1. vB_Kpn_K1PH164C1 2. vB_Kpn_K1lambda5 3. K2PH164C1 4. vB_Kpn_K2PH164C2 5. vB_Kpn_K2alpha62 6. vB_Kpn_K3PH164 7. vB_Ko_K4PH164 8. vB_Ko_K5lambda5 9. vB_Kpn_K6PH25C3 10. vB_Kpn_K7PH164C2 11. vB_Kpn_K7PH164C3 12. vB_Kpn_K7PH164C4 13. vB_Kpl_K8PH128 14. vB_Kpn_K9PH25C2 15. vB_Kpn_K10PH82C1 16. vB_Kpn_K11PH164C1 17. vB_Kpn_K12P1.1 18. vB_Kpn_K13PH07C1L 19. vB_Kpn_K13PH07C1S 20. vB_Kpl_K14PH164C1 21. vB_Kpn_K15PH90 22. vB_Kpn_K16PH164C3 23. vB_Kpn_K17alpha61 24. vB_Kpn_K17alpha62 25. vB_Kpn_K18PH07C1 26. vB_Kpn_K19PH14C4P1 27. vB_Kpn_K2064PH2 28. <colcnt=6> vB_Kpn_K2069PH1 29. vB_Kpn_K2069PH2 30. vB_Kpn_K21lambda1 31. vB_Kpn_K22PH164C1 32. vB_Kpn_K23PH08C2 33. vB_Kpn_K24PH164C1 34. vB_Kpn_K25PH129C1 35. vB_Ko_K26PH128C1 36. vB_Kpn_K27PH129C1 37. vB_Kpn_K28PH129 38. vB_Ko_K29PH164C1 39. vB_Kpn_K30lambda2.2 40. vB_Kpn_K31PH164 41. vB_Kpl_K32PH164C1 42. vB_Kpn_K33PH14C2 43. vB_Kpn_K34PH164 44. vB_Kpl_K35PH164C3 45. vB_Kpn_K36PH128C2B 46. vB_Kpn_K37PH164C1 47. vB_Kpn_K38PH09C2 48. vB_Kpl_K39PH122C2 49. vB_Kpn_K40PH129C1 50. vB_Ko_K41P2 51. vB_Kpn_K42PH8 52. vB_Kpn_K43PH164C1 53. vB_Kpl_K44PH129C1 54. vB_Kpn_K45PH128C2 55. vB_Kpn_K46PH129 56. vB_Kpn_K47PH6 57. vB_Kpl_K48PH164C1 58. vB_Kpl_K49PH164C2 59. vB_Kpn_K50PH164C1 60. vB_Kpn_K51PH129C1 61. vB_Kpn_K52PH129C1 62. vB_Kpl_K53PH164C2 63. vB_Kpl_K54lambda1.1.1 64. vB_Kpn_K54lambda2 65. vB_Kpn_K55P2.1 66. vB_Kpl_K56PH164C1 67. vB_Kpl_K57alpha1.2 68. vB_Kpl_K58PH129C2 69. vB_Kpl_K59PH2 70. vB_Kpn_K60PH164C1 71. vB_Kpn_K61PH164C1 72. vB_Kpn_K62PH164C2 73. vB_Kpn_K63PH128 74. vB_Kpn_K64PH164C4 75. vB_Kte_K65PH164 76. vB_Ko_K66PH128C1 77. vB_Kpn_K68QB 78. vB_Kpn_K69PH164C2 79. vB_Ko_K70PH128C1 80. vB_Kpl_K71PH129C1 81. vB_Kpl_K72PH164C2 82. vB_Ko_K74PH129C2 83. vB_Kpl_K79PH164C1 84. vB_Kpn_K80P2 85. vB_Kpn_K81P1 86. vB_Kpn_K82P1.

Supplementary material 7. **Expected and observed host range of the phage cocktail against the 77 *Klebsiella* spp. reference strains.** (A) Comparative between interactions observed in the spot test of the phage cocktail over each reference strain (O columns) and expected interactions considering the sum of each single phage host range (E columns). Discrepancies are highlighted in black if they are negative and gray if positive (D columns). (B) Comparative between observed and expected cocktail host range over the 77 *Klebsiella* spp. reference strains.

Supplementary material 8. **Anti-phage defense systems profile of the *K. pneumoniae* clinical isolates.** Anti-phage defense system subtypes identified in each clinical isolate.

Supplementary material 9. **Statistical association between ST, KL, OL, AMR genes and susceptibility to the phage cocktail in the panel of clinical isolates**. Statistical tests employed and p-values are specified in the table.

Supplementary material 10. **Role of anti-phage defense systems in resistance to the cocktail of the *K. pneumoniae* clinical isolates.** Each row represents each capsular type with at least one isolate infected by the cocktail. Each column represents a different resistant mechanism. The value of each square represents the differential probability of resistance (dPR) as the difference between the number of isolates with the defense system (DF) resistant to the cocktail (R) and the number of isolates with the defense system susceptible to the cocktail (S). K-types with only one isolate available in de collection, as well as K-types which all isolates are resistant to the cocktail are not included in this figure. Values near to 1 may indicate a relation between having a defense system and an increased resistance to the cocktail.

Supplementary material 11. ***Klebsiella* phages isolated in clinical *K. pneumoniae* and functional annotation.** Genome length, sequencing depth, GC content and taxonomy of the isolated and sequenced *Klebsiella* phages. Annotation of the *Klebsiella* phages isolated in clinical *K. pneumoniae* high-risk clones circulating in Spanish hospitals.

## Notes

### Competing Interest Statement

The authors have declared no competing interest.

