## Supplemental material for "Targeted phage hunting to specific *Klebsiella pneumoniae* clinical isolates is an efficient antibiotic resistance and infection control strategy"

### Supplementary Materials

Supplementary material 1**. Description of the panel of carbapenem-resistant *K. pneumoniae* clinical isolates.** Name and code of the isolates, source, sample type, KL-type, ST, carbapenemase-producing genes and susceptibility to the phage cocktail.

Supp_mat_1.xlsx

Supplementary material 2**. Isolated and sequenced *Klebsiella* phages.** Genome length, sequencing depth, GC content, number of CDS identified, taxonomy, source, accession number of the isolated and sequenced *Klebsiella* phages and list of susceptible KL types.

Supp_mat_2.xlsx

Supplementary material 3. **Functional annotation of *Klebsiella* phages.** Protein-coding gene prediction source for each CDS, start and end position of each one, the DNA strand location and the functional annotation of the sequenced *Klebsiella* phages.

Supp_mat_3.xlsx

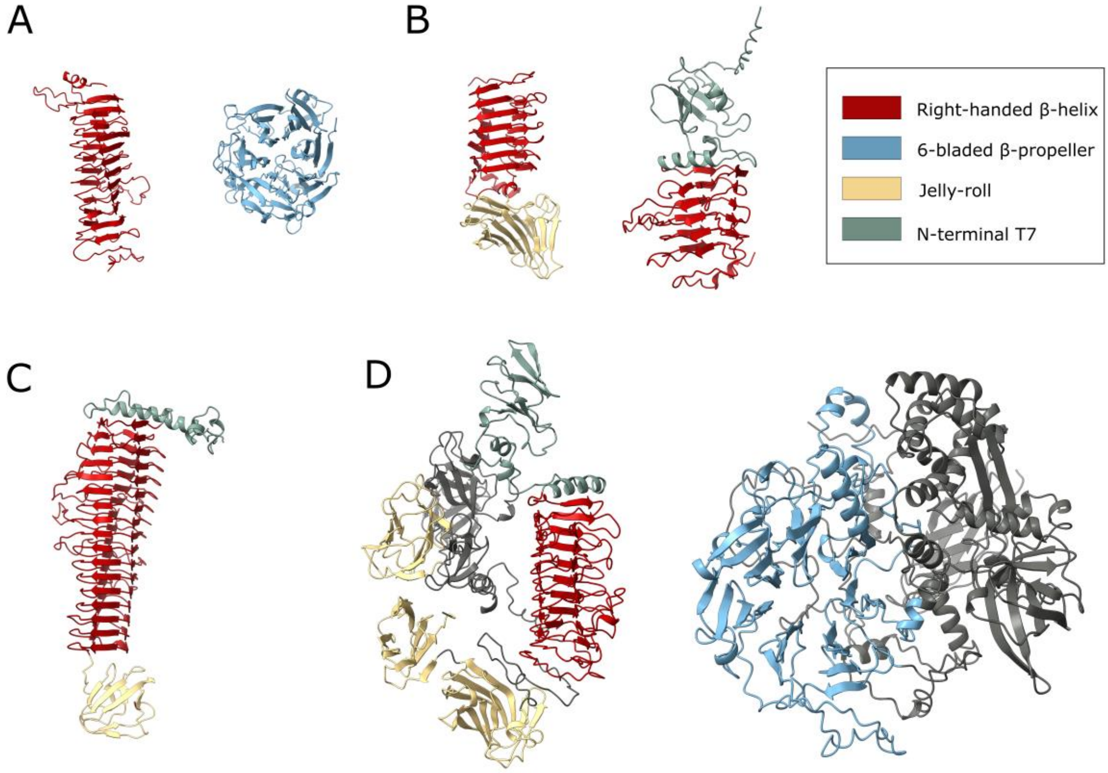


Supplementary material 4. **Architectures of the identified tail spike proteins in the isolated *Klebsiella* phages.** 3D representation of the detected architectures of the tail spike proteins in the isolated phages. (A) The single domain tail spikes. (B) The two domains of tail spikes. (C) the three domains of tail spikes. (D) The complex identified tail spikes. In red the right-handed β-helix, in blue the 6-bladed β-propeller, in gold the jelly-roll, in green the N-terminal domain T7.

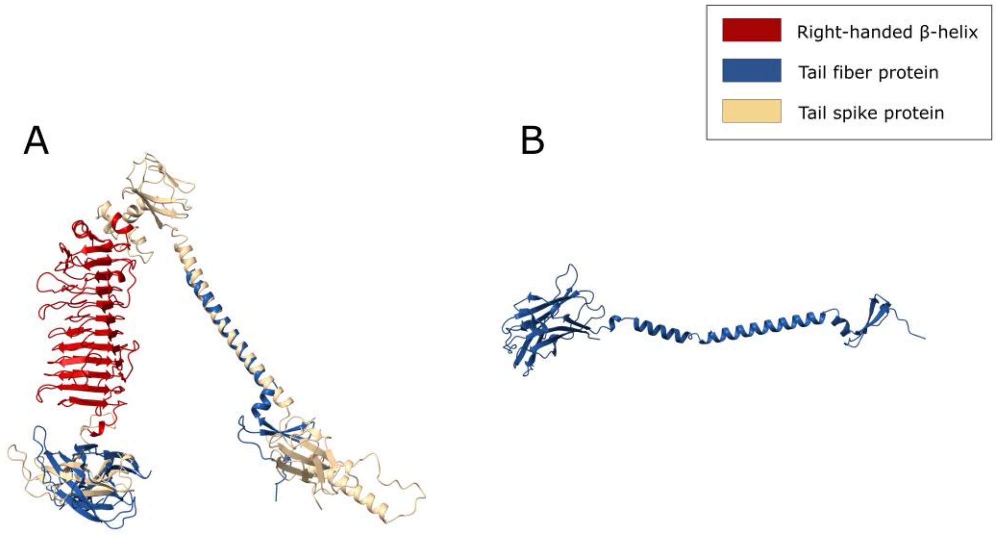


Supplementary material 5. **Illustration of the structural continuity between the tail fiber and the tail spike proteins.** 3D structure representations and alignments of RBPs identified in the isolated phages. In red: the right-handed beta-helix, in blue: the tail fiber protein; in gold: the tail spike protein. (A) Alignment of two protein units (protein unit 1: 1-76; protein unit 2: 77-250) of the tail fiber vB_Kpn_K18PH07C1, CDS 246 and the tail spike vB_Kpl_K39PH122C2, CDS 55 using the Needleman–Wunsch algorithm under Chimerax (85). (B) 3D representation of the tail fiber vB_Kpn_K18PH07C1, CDS 246.

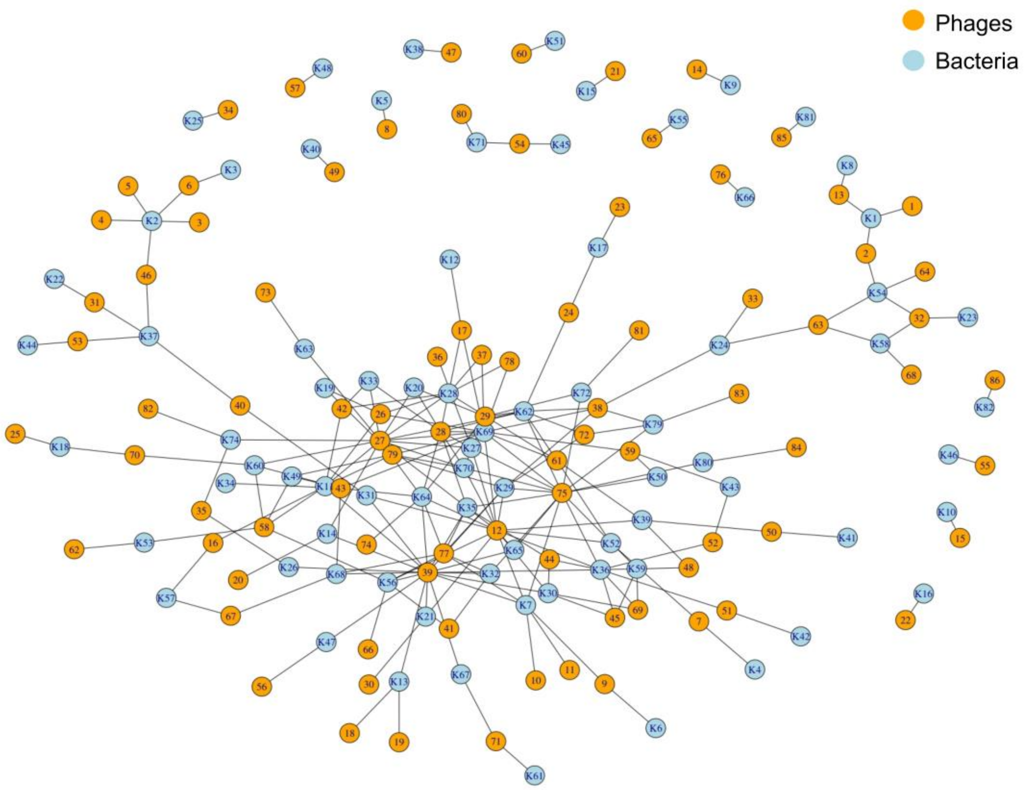


Supplementary material 6. **Network representation of the crossed infection matrix.** We observe 17 disjoint components. Blue nodes represent hosts and orange nodes represent phages. Numbers in orange nodes correspond to these phages: 1. vB_Kpn_K1PH164C1 2. vB_Kpn_K1lambda5 3. K2PH164C1 4. vB_Kpn_K2PH164C2 5. vB_Kpn_K2alpha62 6. vB_Kpn_K3PH164 7. vB_Ko_K4PH164 8. vB_Ko_K5lambda5 9. vB_Kpn_K6PH25C3 10. vB_Kpn_K7PH164C2 11. vB_Kpn_K7PH164C3 12. vB_Kpn_K7PH164C4 13. vB_Kpl_K8PH128 14. vB_Kpn_K9PH25C2 15. vB_Kpn_K10PH82C1 16. vB_Kpn_K11PH164C1 17. vB_Kpn_K12P1.1 18. vB_Kpn_K13PH07C1L 19. vB_Kpn_K13PH07C1S 20. vB_Kpl_K14PH164C1 21. vB_Kpn_K15PH90 22. vB_Kpn_K16PH164C3 23. vB_Kpn_K17alpha61 24. vB_Kpn_K17alpha62 25. vB_Kpn_K18PH07C1 26. vB_Kpn_K19PH14C4P1 27. vB_Kpn_K2064PH2 28. vB_Kpn_K2069PH1 29. vB_Kpn_K2069PH2 30.

vB_Kpn_K21lambda1 31. vB_Kpn_K22PH164C1 32. vB_Kpn_K23PH08C2 33. vB_Kpn_K24PH164C1 34. vB_Kpn_K25PH129C1 35. vB_Ko_K26PH128C1 36. vB_Kpn_K27PH129C1 37. vB_Kpn_K28PH129 38. vB_Ko_K29PH164C1 39. vB_Kpn_K30lambda2.2 40. vB_Kpn_K31PH164 41. vB_Kpl_K32PH164C1 42. vB_Kpn_K33PH14C2 43. vB_Kpn_K34PH164 44. vB_Kpl_K35PH164C3 45. vB_Kpn_K36PH128C2B 46. vB_Kpn_K37PH164C1 47. vB_Kpn_K38PH09C2 48. vB_Kpl_K39PH122C2 49. vB_Kpn_K40PH129C1 50. vB_Ko_K41P2 51. vB_Kpn_K42PH8 52. vB_Kpn_K43PH164C1 53. vB_Kpl_K44PH129C1 54. vB_Kpn_K45PH128C2 55. vB_Kpn_K46PH129 56. vB_Kpn_K47PH6 57. vB_Kpl_K48PH164C1 58. vB_Kpl_K49PH164C2 59. vB_Kpn_K50PH164C1 60. vB_Kpn_K51PH129C1 61. vB_Kpn_K52PH129C1 62. vB_Kpl_K53PH164C2 63. vB_Kpl_K54lambda1.1.1 64. vB_Kpn_K54lambda2 65. vB_Kpn_K55P2.1 66. vB_Kpl_K56PH164C1 67. vB_Kpl_K57alpha1.2 68. vB_Kpl_K58PH129C2 69. vB_Kpl_K59PH2 70. vB_Kpn_K60PH164C1 71. vB_Kpn_K61PH164C1 72. vB_Kpn_K62PH164C2 73. vB_Kpn_K63PH128 74. vB_Kpn_K64PH164C4 75. vB_Kte_K65PH164 76. vB_Ko_K66PH128C1 77. vB_Kpn_K68QB 78. vB_Kpn_K69PH164C2 79. vB_Ko_K70PH128C1 80. vB_Kpl_K71PH129C1 81. vB_Kpl_K72PH164C2 82. vB_Ko_K74PH129C2 83. vB_Kpl_K79PH164C1 84. vB_Kpn_K80P2 85. vB_Kpn_K81P1 86. vB_Kpn_K82P1.

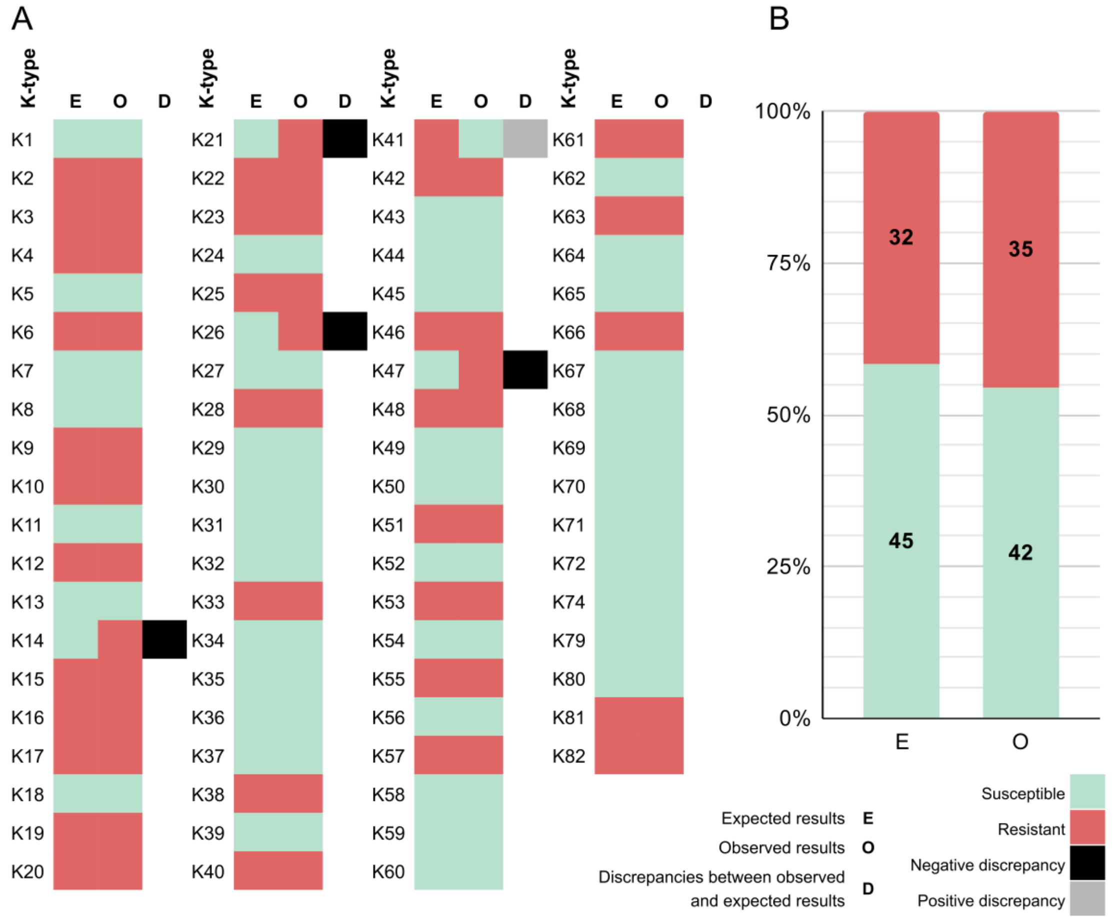


Supplementary material 7. **Expected and observed host range of the phage cocktail against the 77 *Klebsiella* spp. reference strains.** (A) Comparative between interactions observed in the spot test of the phage cocktail over each reference strain (O columns) and expected interactions considering the sum of each single phage host range (E columns). Discrepancies are highlighted in black if they are negative and gray if positive (D columns). (B) Comparative between observed and expected cocktail host range over the 77 *Klebsiella* spp. reference strains.

Supplementary material 8. **Anti-phage defense systems profile of the *K. pneumoniae* clinical isolates.** Anti-phage defense system subtypes identified in each clinical isolate.

Supp_mat_8.xlsx

Supplementary material 9. **Statistical association between ST, KL, OL, AMR genes and susceptibility to the phage cocktail in the panel of clinical isolates**. Statistical tests employed and p-values are specified in the table.

Supp_mat_9.xlsx

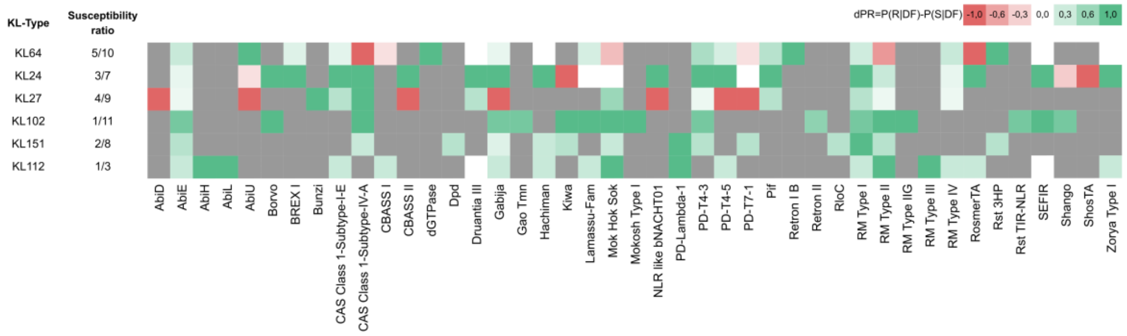


Supplementary material 10. **Role of anti-phage defense systems in resistance to the cocktail of the *K. pneumoniae* clinical isolates.** Each row represents each capsular type with at least one isolate infected by the cocktail. Each column represents a different resistant mechanism. The value of each square represents the differential probability of resistance (dPR) as the difference between the number of isolates with the defense system (DF) resistant to the cocktail (R) and the number of isolates with the defense system susceptible to the cocktail (S). K-types with only one isolate available in de collection, as well as K-types which all isolates are resistant to the cocktail are not included in this figure. Values near to 1 may indicate a relation between having a defense system and an increased resistance to the cocktail.

Supplementary material 11. ***Klebsiella* phages isolated in clinical *K. pneumoniae* and functional annotation.** Genome length, sequencing depth, GC content and taxonomy of the isolated and sequenced *Klebsiella* phages. Annotation of the *Klebsiella* phages isolated in clinical *K. pneumoniae* high-risk clones circulating in Spanish hospitals.

Supp_mat_11.xlsx
